# Quantitative multi-locus metabarcoding and waggle dance interpretation reveal honey bee spring foraging patterns in Midwest agroecosystems

**DOI:** 10.1101/418590

**Authors:** Rodney T. Richardson, Hailey R. Curtis, Emma G. Matcham, Chia Hua-Lin, Sreelakshmi Suresh, Douglas B. Sponsler, Luke E. Hearon, Reed M. Johnson

## Abstract

We explored the pollen foraging behavior of honey bee colonies situated in the corn and soybean dominated agroecosystems of central Ohio over a month-long period using both pollen metabarcoding and waggle dance inference of spatial foraging patterns. For molecular pollen analysis we developed simple and cost-effective laboratory and bioinformatics methods. Targeting four plant barcode loci (ITS2, *rbcL, trnL* and *trnH*), we implemented metabarcoding library preparation and dual-indexing protocols designed to minimize amplification biases and index mis-tagging events. We constructed comprehensive, curated reference databases for hierarchical taxonomic classification of metabarcoding data and used these databases to train the Metaxa2 DNA sequence classifier. Comparisons between morphological and molecular palynology provide strong support for the quantitative potential of multi-locus metabarcoding. Results revealed consistent foraging habits between locations and show clear trends in the phenological progression of honey bee spring foraging in these agricultural areas. Our data suggest that three key taxa, woody Rosaceae such as pome fruits and hawthorns, *Salix*, and *Trifolium* provided the majority of pollen nutrition during the study. Spatially, these foraging patterns were associated with a significant preference for forests and tree lines relative to crop fields and herbaceous land cover.

## Introduction

Understanding the floral resource usage patterns and preferences of pollinators, such as honey bees, remains an important research goal with implications for pollinator health (Di Pasquale et al. 2013; Vaudo et al. 2016). Such questions have typically been investigated using plant-pollinator network analysis and analysis of pollen provisions (Severson and Parry 1981; Memmott 1999). In the case of honey bees, however, waggle dance inference of spatial foraging patterns has emerged as an additional tool for investigating the relative attractiveness of different landscape features as forage and inferring associations between landscape composition and foraging outcomes (Couvillon and Ratnieks 2015). In this study, we combined molecular pollen analysis methods with waggle dance inference to observe the taxonomic composition of honey bee-collected pollen while simultaneously inferring where bees were foraging in the surrounding landscape.

Since the first proof-of-concept articles documenting the applicability of plant metabarcoding to pollen analysis (Valentini et al. 2010; Hawkins et al. 2015; Keller et al. 2015; Kraaijeveld et al. 2015; Richardson et al. 2015), the field of molecular pollen analysis has expanded rapidly (Cornman et al. 2016; McFrederick and Rehan 2016; Smart et al. 2017; Bell et al. 2018). Undoubtedly, high-throughput sequencing exhibits great promise in facilitating future discoveries in the fields of plant-pollinator interaction biology, palynological forensics, food authentication and airborne pollen monitoring (Bell at al. 2016). Despite this promise, important questions remain regarding the selection of appropriate library preparation protocols and bioinformatic analysis methods. Further, the ability to draw quantitative inferences from pollen metabarcoding studies remains unclear, with considerable disagreement between research groups (Keller et al. 2015; Richardson et al. 2015; Bell et al. 2018).

For any researcher wishing to employ pollen metabarcoding, the selection of library preparation protocols is a critical methodological decision. The selection of which loci to target, which universal primers to use for amplification and which library construction methods to implement will ultimately affect the strengths or weaknesses of the study, regardless of the bioinformatic techniques employed after sequencing. With respect to locus and primer choice, a number of studies have documented the taxonomic biases of different primer sets used to amplify the same locus (Deagle et al. 2014; Elbrecht and Leese 2015; Piñol et al. 2015; Krehenwinkel et al. 2017), as well as the biases of individual loci (Cowart et al. 2015; Richardson et al. 2015a; Elbrecht et al. 2016). Similarly, it has recently been demonstrated that the use of ‘barcoded’ or ‘fusion’ primers during the initial amplification of mixed-species samples results in considerable amplification bias and decreased replicability (Berry et al. 2011; O’Donnell et al. 2016). While such primers are used to attach oligonucleotides necessary for indexing and next-generation sequencing, both studies demonstrated that this could be performed with greater replicability and precision by first performing a traditional PCR with no 5’ appendages on the primer before using a second set of fusion primers carrying nucleotide sequences required for sequencing. It is likely that biases resulting from sub-optimal primer and locus selection combined with the use of fusion primers for initial community amplification will result in sequencing data that does not quantitatively represent the diversity of pollen being analyzed.

Given the above issues, the use of multi-locus metabarcoding has been proposed as one approach to improving the quantitative capacity of molecular pollen analysis. Since different loci and primer sets display different biases with respect to the taxonomic scope of detection and quantitative bias, employing multiple markers and analyzing the median or mean of all loci may improve the accuracy of quantitative inferences (Richardson et al. 2015a). This approach has the added advantage of enabling researchers to exclude taxa identified using only one locus and focus on consensus taxa identified by multiple markers, increasing the confidence of detections.

Following sequencing, the bioinformatic characterization of the resulting data is another important consideration. Arguably, metabarcoding studies should include rigorous tests of DNA sequence classification methods to enable reviewers and readers to properly gauge the plausibility of research findings (Edgar 2018). This requires tests of both the accuracy and sensitivity of bioinformatics methods and leads researchers toward methods that can be benchmarked against alternative approaches. While this requires extra effort, it enables researchers to be more objective in selecting classification methods and to rely less on *post hoc* determination of classification parameters while working through preliminary analyses of their data.

To classify our pollen metabarcoding data, we employed a recently designed DNA sequence classifier, Metaxa2 (Bengtsson-Palme et al. 2015). The Metaxa2 classifier is capable of extracting sequences belonging to a specific locus of interest from multi-locus or metagenomic data using Hidden Markov Models produced by HMMER (Eddy 2011). Since Metaxa2 had not previously been trained on plant barcode loci, we produced curated plant reference databases for each of our loci of interest and performed a cross-validation analysis as in Richardson et al. (2017). Using logistic regression, we characterized the relationship between the Metaxa2 reliability score and the probability of false classification. We then used this regression model to select a classification reliability score threshold optimized for our data and reference databases. Finally, we examined the accuracy and sensitivity of Metaxa2 implemented with our chosen reliability score threshold using previously documented methods (Richardson et al. 2018).

An overarching goal of this work was to develop complementary laboratory and bioinformatics approaches optimized to minimize quantitative biases in a cost-effective manner. Using a three-step PCR approach, we circumvent the issue of using fusion primers in the initial sample amplification, similar to the approach suggested in Berry et al. (2011) and O’Donnell et al. (2016). Further, due to the presence of critical mis-tag events in next-generation sequencing data (Schnell et al. 2015), we performed our experiment using a 50 percent unsaturated Latin Square Design, as described in Esling et al. (2015). To accomplish this efficiently, we utilized the gene annotation capacity of Metaxa2 to minimize the number of dual index pairs required for our study. This allowed us to produce multiple libraries per sample using the same dual index pair and computationally separate sequences from each locus after sequencing. Sequencing multiple loci per sample on the same Illumina flow cell has the added advantage of increasing sequence diversity during initial base calling, decreasing the amount of PhiX required in the final amplicon pool and increasing the number of samples which can be analyzed per sequencing run.

While methods will continue to be optimized, especially with respect to locus choice, primer choice and bioinformatic classification, our work represents a simplified and cost-effective approach to pollen metabarcoding which yields quantitatively useful data. Further, we demonstrate the applicability of our methods through applying them to explore the foraging ecology of honey bee, *Apis mellifera*, colonies situated across four apiaries in the corn and soybean agroecosystems of west-central Ohio.

In conducting a waggle dance analysis study in tandem with our pollen metabarcoding approach, we were able to relate the taxonomic composition of our samples to observed spatial patterns of honey bee foraging, as in Sponsler et al. (2017). Waggle dance inference was useful for determining the relative importance of different landclass types as honey bee foraging locations for our study. However, the relatively high degree of imprecision inherent in this type of analysis made fine-scale interpretation of foraging patterns unfeasible and resulted in low statistical power with respect to inferring differences in foraging preference across landscape classes despite considerable sampling effort.

## Methods

### Pollen sampling and waggle dance recording

In early spring of 2015, apiaries were set up at four sites in rural central Ohio. Each apiary consisted of 12 - 18 actively foraging colonies in 8- or 10-frame Langstroth hives. Two of the Langstroth hives were fitted with Sundance I bottom-mounted pollen traps (Ross Rounds, Albany, NY, USA). Pollen was trapped continuously from May 2^nd^ to May 27^th^. The traps were emptied and samples were collected at three to five day intervals. Artificial pollen substitute (Ultra Bee, Mann Lake, Hackensack, MN, USA) were placed in the pollen-trapping hives to mitigate the effects of the resulting pollen nutritional deficit. To video record waggle dancing behavior, one 3-frame observation hive (Bonterra TableView, Addison, ME, USA) was installed at each apiary, sheltered in a plastic storage shed (Suncast #BMS4700, 179×112×132 cm, Batavia, IL, USA). Approximately one hour of video of the bottom frame was recorded using an HD video camera (Canon Vixia HF G20) situated on a 1m tripod with light provided by a small opening in the door. Recordings were made on 16 days from May 4^th^ to May 29^th^ between the hours of 10:30 and 17:10. In total, video recordings were taken on at least seven sampling dates per apiary throughout the study.

### Metabarcoding sample processing

For each sample, 10 percent by mass or up to 20 g of pollen (wet mass) was combined with distilled water to a concentration of 0.1 g/mL of pollen and homogenized using a blender (Hamilton Beach #54225, Southern Pines, NC, USA) for 2.5 minutes. After blending, each sample was gently mixed immediately prior to the collection of 1.4 mL of pollen homogenate into a 2.0 mL bead beater tube (Fisherbrand Free-Standing Microcentrifuge Tubes; Fisher Scientific, Hampton, NH, USA). Bead beater tubes were then centrifuged for 2 minutes at 10,000 g, the supernatant was removed from the pollen pellet and 1.25 mL of buffer AP1 from the Qiagen DNeasy Plant Minikit (QIAGEN, Venlo, The Netherlands) was added along with 3,355 mg of 0.7 mm zirconia beads (Fisher Scientific, Hampton, NH, USA). Pollen was then mechanically disrupted in a bead beater (Mini-BeadBeater-1; BioSpec Products, Bartlesville, OK, USA) for 5 minutes at the highest setting. Samples were then vortexed briefly before 400 µL of lysate was removed for DNA extraction using the Qiagen DNeasy Plant Minikit according to the manufacturer’s instructions.

Following DNA extraction, a 3-step PCR-based protocol, compatible with the Illumina Nextera sequencing protocol, was used for amplicon library preparation. For each sample, *rbcL, trnL, trnH* and ITS2 libraries were prepared separately using previously published universal primer sets (White et al. 1990; Fay et al. 1997; Sang et al. 1997; Tate and Simpson 2003; Taberlet et al. 2007; Chen et al. 2010). For the initial PCR reaction, universal primers with no 5-prime fusion oligos were used to generate a pool of amplicons. Subsequently, 1 µL of PCR product from the initial reaction was used as template for a second PCR reaction. Lastly, 1 µL of PCR product from the second reaction was used as template for a third PCR reaction. The second and third reactions were used to append template priming, sample indexing and lane hybridization oligonucleotides to each amplicon for downstream compatibility with the Illumina Nextera protocol and MiSeq sequencing. Supplementary Table S1 contains the primer sequences, complete PCR conditions and sample dual-indexing design used in this study. All PCR reactions were conducted at a 20 μl scale with 4 μl High Fidelity Phusion Buffer, 0.2 mM dNTPs and 0.02 U/μl Phusion Polymerase. Initial PCR reactions were conducted with 100 to 150 ng of DNA template. Following library preparation, the final PCR products were purified and normalized using the SequalPrep Normalization Plate Kit (Thermo Fisher Scientific, Waltham, MA, USA), pooled equimolarly and sequenced using the Illumina MiSeq Micro Kit (2 x 150 cycles).

### Hierarchical classification database construction and curation

To use Metaxa2 (v2.2 4^th^ beta; Bengtsson-Palme et al. 2015), a software originally designed to classify bacterial and fungal sequences, for plant sequence classification, we first had to produce reference databases for each marker of interest. To gather reference data, we downloaded all available *trnL, trnH, rbcL* and plant whole chloroplast genome sequences from NCBI Genbank on April 20^th^, 2017. Additionally, we downloaded all available Viridiplantae ITS2 sequences from the ITS2 Database (Ankenbrand et al. 2015) on May 5^th^, 2017. We then used the NCBI Taxonomy Module (NCBI Resource Coordinators, 2018) along with the Perl scripts described in Sickel et al. (2015) to obtain the seven-ranked Linnaean lineage, from kingdom to species, for each reference entry.

To aid in both the estimation of marker conservation parameters during hierarchical training and the removal of duplicate reference sequences, we extracted the amplicon of interest from the available reference sequences where possible, including from plant whole chloroplast genomes for the *rbcL* and *trnL* markers. For this, we first removed exceptionally long or short sequence entries as well as any entries containing three or more consecutive uncalled base pairs from the locus-specific data sets. For the *rbcL, trnL* and whole chloroplast genome data sets, we then isolated archetypical *trnL* and *rbcL* reference sequences for each locus using the primers employed during pollen metabarcoding and used these sequences in combination with the HMM-based Metaxa2 Database Builder tool (v1.0 4^th^ beta; Bengtsson-Palme et al. 2018) to extract the amplicon of interest. However, *trnH* sequences were too divergent for this approach. Instead, we removed any entries longer than 1,500 bp and retained only the references annotated with ‘*trnH’* and ‘*psbA’*, to remove as many extraneous sequences as possible. While ITS2 is also a highly divergent marker, this level of curation for ITS2 references was unnecessary due to the *in silico* secondary structure analysis employed during ITS2 Database curation (Keller et al. 2009).

Next, we performed extensive curation of the taxonomic lineage metadata associated with each entry. Using Perl substitution, we removed the undefined ranks from the end of any lineage unidentified at the highest resolution ranks, typically genus and species. Leaving undefined tags in the lineages is problematic for hierarchical classification, as the classifier has no way to distinguish undefined annotations from *bona fide* taxonomic annotations, resulting in multiple sequences from different taxa receiving the same annotation. To account for lineages which are currently unresolved at intermediate ranks, we developed a Python script which substitutes these undefined intermediate rank annotations with an annotation containing the identity of the lowest resolution rank containing an identification and a ‘urs’ tag which indicates ‘unresolved.’ In this way, we were able to salvage important lineages of plants, such as Magnoliales, Ranunculales and Caryophyllales, while annotating these entries with a tag that distinguishes them from other taxa that are also unresolved at the same rank. For a more detailed description of this approach, see Richardson et al. (2018). Finally, we used Perl substitution to further clean the lineages and remove ranks annotated with artifactual alphanumeric tags or open nomenclature, which were common at the genus, species and family ranks. Reference sequence databases were then dereplicated using the Java script provided with the RDP Naïve Bayesian Classifier (v2.11; Wang et al 2007).

Following final curation of the reference sequence and taxonomic lineage data, Metaxa2 was trained on each of the four markers using the Metaxa2 Database Builder Tool. For *rbcL* and *trnL*, training was performed in default mode and an archetypical sequence was used to designate the precise barcode region of interest. For *trnH* and ITS2, the divergent mode was used due to the low degree of sequence conservation across these markers.

In addition to training Metaxa2 on the complete reference databases for each marker, we also performed a cross-validation performance evaluation by randomly sampling 10 percent of the sequences for each marker to serve as test sequences, training Metaxa2 with the remaining 90 percent and then classifying the test sequences. In order to make the evaluation conservative, we cropped the test sequences for each marker to 150 bp in length using a custom Python script. To select the most appropriate Metaxa2 reliability score, an estimate of classification confidence, we evaluated the relationship between the Metaxa2 reliability score and classification error probability using local polynomial logistic regression. For this evaluation, we randomly subsampled 1,000 reference sequence classification cases from each locus and regressed the outcome of each family-level classification case, ‘0’ indicating correct classification and ‘1’ indicating misclassification, against the Metaxa2 reliability score of the assignment using the Loess function in R (R Core Team 2014). We then estimated the sensitivity and accuracy of the classifier using the methods of Richardson et al. (2017).

### Pollen metabarcoding bioinformatics and statistics

Given the built-in quality filtering and mate-pair awareness of Metaxa2, we proceeded to classify the sequences of the raw forward and reverse fastq files without prior quality processing for the ITS2, *trnH* and *rbcL* libraries. Since amplicons of the *trnL* marker were short enough for the paired end reads to be merged into a single contiguous sequence, we used PEAR (v0.9.1; Zhang et al. 2014) to merge forward and reverse read pairs and improve base calling accuracy toward the middle of the *trnL* amplicon. For read pairing, a minimum merged read length of 100 bp was used along with a Phred scale 33 quality threshold of 20. Assembled *trnL* sequences were then subjected to taxonomic classification using Metaxa2. For all taxonomic classification, Metaxa2 was implemented using default quality filtering and a reliability score threshold of 50 on the Owens cluster of the Ohio Supercomputer Center (Ohio Supercomputer Center, 1987).

Following sequence classification, custom Python scripts were used to summarize the data using the consensus-filtered, median-based approach discussed in Richardson et al. (2015a). Briefly, for a given sample and taxonomic rank, the proportion of sequences belonging to each taxon was calculated for each marker. At this point, the taxa were consensus-filtered by discarding any taxonomic group which was discovered using only one of the four makers. Additionally, taxonomic groups represented by less than 0.01 percent of the data were discarded. The median proportional abundance of each taxonomic group was then calculated. After obtaining the median proportions of each taxon, median values were then normalized to a sum of 1.0 for each sample. The commands and Python scripts used for all analyses presented in this work are available at https://github.com/RTRichar/QuantitativePollenMetabarcoding.

### Microscopic palynology and quantitative inference

To explore the utility of metabarcoding data for drawing quantitative inferences of the proportions of different taxa within each sample, we used microscopic palynology as a standard to characterize the components of 12 out of the 32 total pollen samples. We then performed linear regression analysis, regressing the metabarcoding data against microscopic inferences of the abundance of each taxon for each sample analyzed. For these analyses, all regressions were performed using data summarized to the family rank. For microscopic characterizations, we utilized the methods of Richardson et al. (2015b) wherein corbicular pollen pellets were first sorted by color prior to mounting, basic fuchsin staining and microscopic identification. Following the taxonomic characterization of each color fraction, the sum proportion of each taxonomic group was calculated according to the volumetric methods of O’Rourke and Buchmann (1991). The pollen reference collections used for identification were those detailed in Richardson et al. (2015a and b).

### Waggle dance analysis and statistics

Waggle dance analysis was conducted using methods similar to Sponsler et al. (2017). Briefly, each video was subsampled by extracting one-minute segments separated by four minute intervals. Individual dance vectors (distance and direction) were then estimated according to Couvillon et al. (2012) using ImageJ (Schneider et al. 2012) video analysis with the MTrackJ plugin (Meijering et al., 2012). Using QGIS (v2.18.20; QGIS Development Team 2018), we digitized the landscape within a 2 km radius of each apiary using the USDA-NASS Cropland Data Layer (USDA National Agricultural Statistics Service Cropland Data Layer 2018), OpenLayers aerial imagery (Map data provided by Google; Sourcepole 2018), and manual ground-truthing. Landscape features were classified into three categories: crop field, forest (forest and tree lines) and herbaceous habitat (residential and pasture lands). Dance vectors were then mapped upon the digitized landscape using the Bayesian probabilistic methods of Schürch et al. (2013). For each landcover class, we calculated a preference index, defined as the empirical visitation rate on a given landcover class (i.e. the sum of the foraging probability falling within a given landcover class) divided by the proportional abundance of that landcover class in the total landscape. Conceptually, this is a measure of whether the landcover class visitation rate deviates from what would be expected assuming random foraging across the landscape. After applying a log transformation to this statistic, values above zero indicate preference for the landcover class, while values below zero indicate aversion. For statistical analysis, we applied a one-way anova to our log transformed preference index values to infer if significant differences in preference existed across the three landcover types. Additionally, we used two-tailed t-tests to infer if the preference index of any of the three landcover types was significantly different from zero. Lastly, since honey bees prioritize floral resource use based on distance from the hive, we used a one-way ANOVA to test for significant differences in mean distance from the apiary across landcover types.

## Results

### Construction and evaluation of hierarchical classification databases

Construction and curation of reference sequence databases yielded 21,902, 22,663, 46,488 and 121,168 sequences for *rbcL, trnL, trnH* and ITS2, respectively. These sequences corresponded to between 16,994 to 65,052 species per database and a total of 86,525 species across all four databases (Table 1). With respect to classification performance, local polynomial logistic regression between reference test sequence classification outcome and the “reliability score” calculated by Metaxa2 revealed a non-linear relationship when classifying 150 bp plant reference sequences (Figure 1A). The probability of classification error was below 0.1 for reliability scores of 54 or greater. To maximize sensitivity, we chose to set the reliability score at 50 for analysis of all four loci. Using this threshold in our accuracy and sensitivity assessments, we found Metaxa2 to exhibit a low degree of error, mis-identifying an average of 5.1, 2.0 and 1.2 percent of 150 bp plant reference test sequences at the level of genus, family and order, respectively. Further, we found high degrees of sensitivity in classifying 150 bp plant reference test sequences, with an average genus-level sensitivity of 40.4 percent and family and order-level sensitivities of 89.8 and 94.4 percent, respectively (Figure 1B).

**Table 1:** Summary of plant taxonomic groups represented at each rank in the reference sequence databases. Estimates for intermediate nodes of the phylogenetic tree, predominantly class and order, are possibly artificially inflated do to some unresolved lineages from the same taxon being given their own independent annotations for hierarchical classification purposes.

|  | Phyla | Classes | Orders | Families | Genera | Species |
| --- | --- | --- | --- | --- | --- | --- |
| ITS2 | 2 | 53 | 171 | 612 | 9524 | 65052 |
| *rbcL* | 2 | 63 | 196 | 776 | 7000 | 16994 |
| *trnL* | 1 | 30 | 84 | 374 | 5943 | 18169 |
| *trnH* | 1 | 42 | 126 | 517 | 5341 | 26288 |
| **Total** | **2** | **69** | **229** | **879** | **12952** | **86525** |

**Figure 1:**
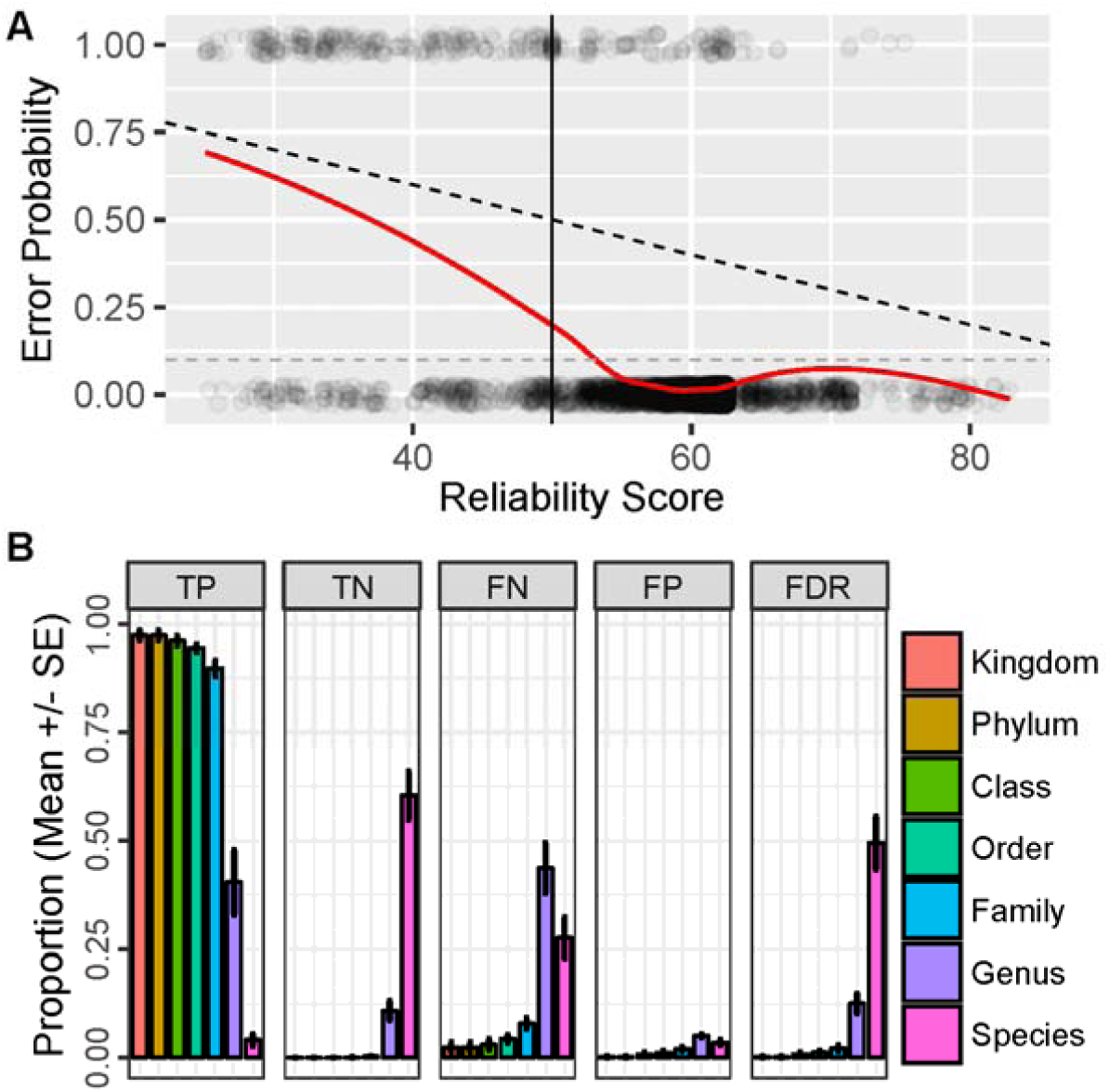
Cross-validation results of classifier performance evaluation on test reference sequences cropped to 150 bp in length. Local polynomial logistic regression of test case classification outcomes, ‘1’ indicating an incorrect classification and ‘0’ indicating a correct classification, regressed against Metaxa2 reliability score (A). A dashed black line illustrates the hypothetically ideal relationship between error probability and the reliability score. The best fit local polynomial model for the data is shown with a solid red line and a dashed grey line indicates an error probability of 0.1. Mean and standard error of the proportion of true positive (TP), true negative (TN), false negative (FN) and false positive (FP) classifications as well as the classification false discovery rate (FDR) across each taxonomic rank for all four plant markers (B).

### Sequencing, demultiplexing and classification performance

After sequencing, we obtained 4,380,260 mate-paired reads. Of these 3,141,670 mate-pairs were classified as Viridiplantae by Metaxa2 using HMM-based sequence annotation and extraction. Following extraction of the sequences from each locus, we obtained a mean and standard error of 25,572 ± 1,416 sequences per sample per locus. An ANOVA followed by a Tukey’s HSD test revealed significant differences in the average number of sequences per sample across the four loci used (ANOVA: *P* < 0.0001; *P* < 0.0001 for all pairwise comparisons except ITS2 - *trnH, P* < 0.05, and ITS2 - *trnL, P* > 0.05). Overall, the minimum number of Viridiplantae sequences found in a single library was 1,897. Table 2 shows the mean, standard error and range of sequences per sample for each locus. Following sequence classification with Metaxa2, we generally achieved a high rate of classification from phylum to family, beyond which, steep decreases in sensitivity were observed at the genus and species ranks. However, one marker, ITS2, exhibited relatively high sensitivity at the genus level. This was expected given the increased discriminatory power of ITS2 relative to other plant barcodes (Chen et al. 2010). For sequences belonging within Viridiplantae, Table 3 shows the mean proportion of sequences classified and standard error for each marker at each taxonomic rank.

**Table 2:**
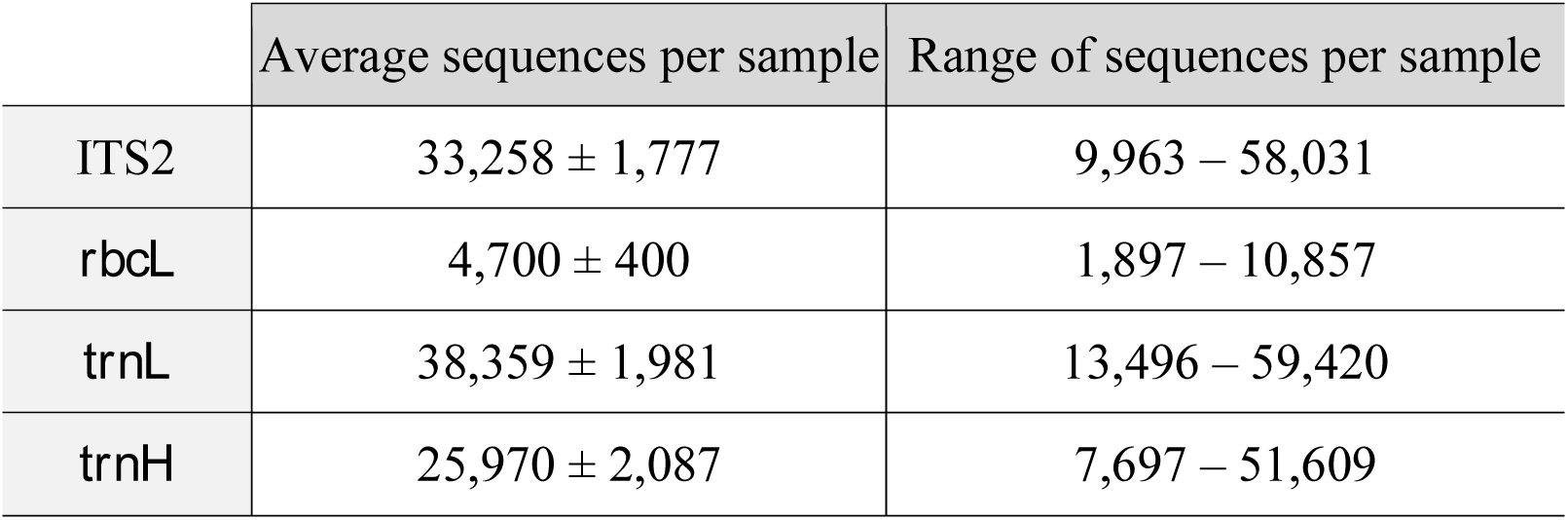
Mean, standard error and range of the number of Viridiplantae sequences per sample obtained for each marker.

|  | Average sequences per sample | Range of sequences per sample |
| --- | --- | --- |
| ITS2 | 33,258 ± 1,777 | 9,963 – 58,031 |
| *rbcL* | 4,700 ± 400 | 1,897 – 10,857 |
| *trnL* | 38,359 ± 1,981 | 13,496 – 59,420 |
| *trnH* | 25,970 ± 2,087 | 7,697 – 51,609 |

**Table 3:** Mean and standard error of proportion of sequences classified to each rank for each marker.

|  | ITS2 | *rbcL* | *trnL* | *trnH* |
| --- | --- | --- | --- | --- |
| Kingdom | 0.89 ± 0.02 | 1.00 ± 0.00 | 1.00 ± 0.00 | 0.99 ± 0.00 |
| Phylum | 0.89 ± 0.02 | 1.00 ± 0.00 | 1.00 ± 0.00 | 0.99 ± 0.00 |
| Class | 0.83 ± 0.03 | 0.96 ± 0.01 | 0.99 ± 0.00 | 0.99 ± 0.00 |
| Order | 0.75 ± 0.03 | 0.87 ± 0.01 | 0.98 ± 0.00 | 0.99 ± 0.00 |
| Family | 0.70 ± 0.03 | 0.78 ± 0.03 | 0.97 ± 0.00 | 0.99 ± 0.00 |
| Genus | 0.27 ± 0.03 | 0.15 ± 0.02 | 0.23 ± 0.03 | 0.47 ± 0.04 |
| Species | 0.03 ± 0.01 | 0.03 ± 0.01 | 0.10 ± 0.02 | 0.11 ± 0.03 |

### Quantitative median-based multi-locus metabarcoding

With respect to the quantitative validity of metabarcoding data, we found extreme variance in the degree to which results from different loci were related to the microscopic results using linear regression modeling (Figure 2). Prior to these analyses, the microscopy and molecular datasets were square-root transformed in order to improve homogeneity of variance, which is negatively affected by the preference of honey bees to collect small quantities of numerous plant taxa. When using our multi-locus approach, we found the metabarcoding median of each consensus-filtered family to be strongly and significantly related to the microscopy results (*P* < 0.0001; *R*^2^ = 0.60). Analyzing individual loci, the results from *rbcL* and *trnL* were strongly correlated with the microscopy results (*P* < 0.0001 and *R*^2^ > 0.53 for both loci). Further, while the *trnH* results were significantly correlated with the microscopy results, this relationship was relatively weak (*P* < 0.0001; *R*^2^ = 0.31). Lastly, the data from the ITS2 locus was not significantly related to the microscopy results (*P* > 0.05; *R*^2^ = −0.001).

**Figure 2:**
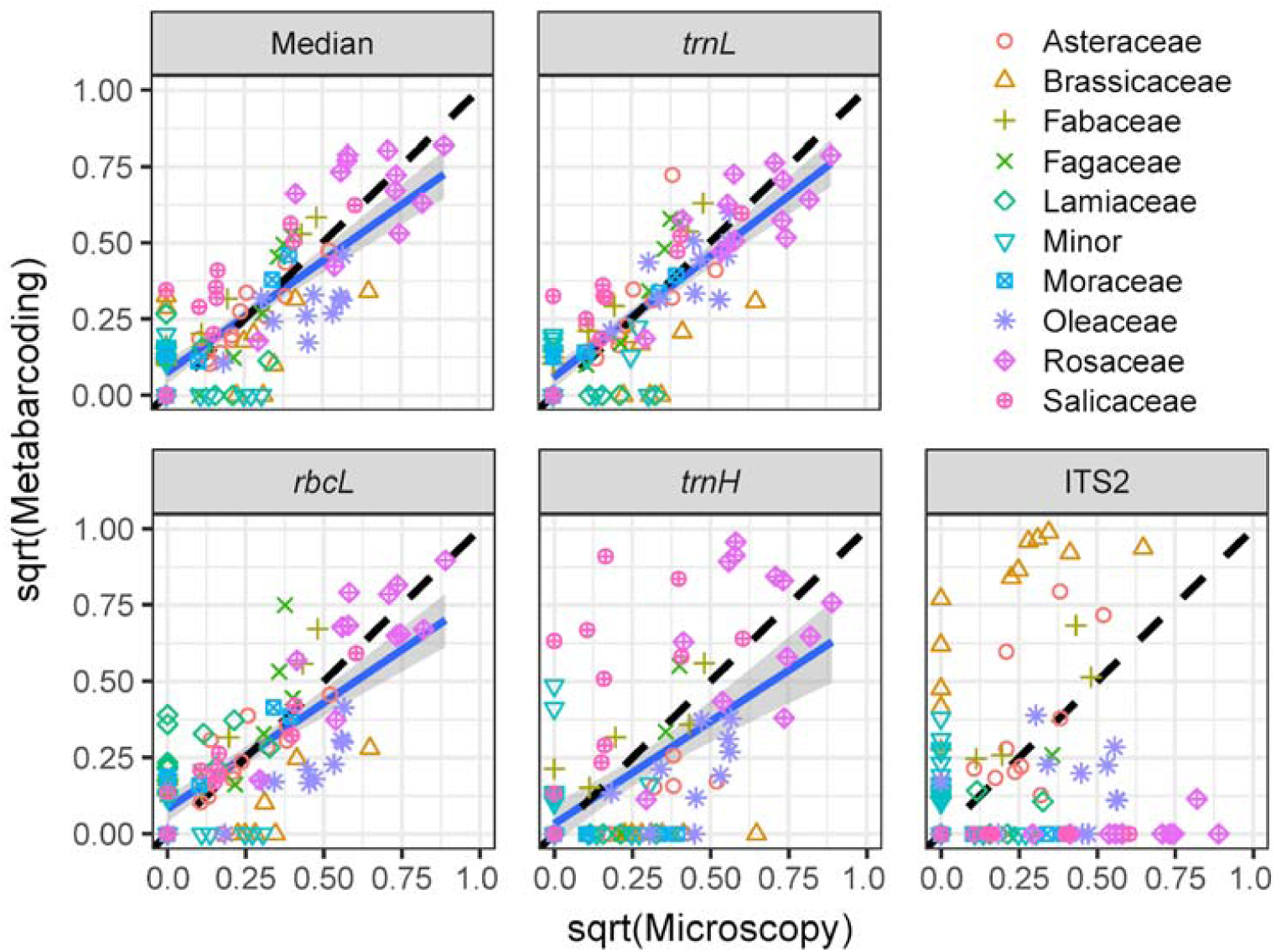
Metabarcoding results regressed against microscopy results for the metabarcoding median of all loci as well as each locus individually. All proportional results are summarized to the family level and proportions are square-root transformed. Plant families occurring in any sample at greater than 5 percent in the metabarcoding median results are shown with distinct colors and point types and both the molecular and microscopic results were filtered to remove detections of less than 1 percent of the untransformed data.

**Figure 3:**
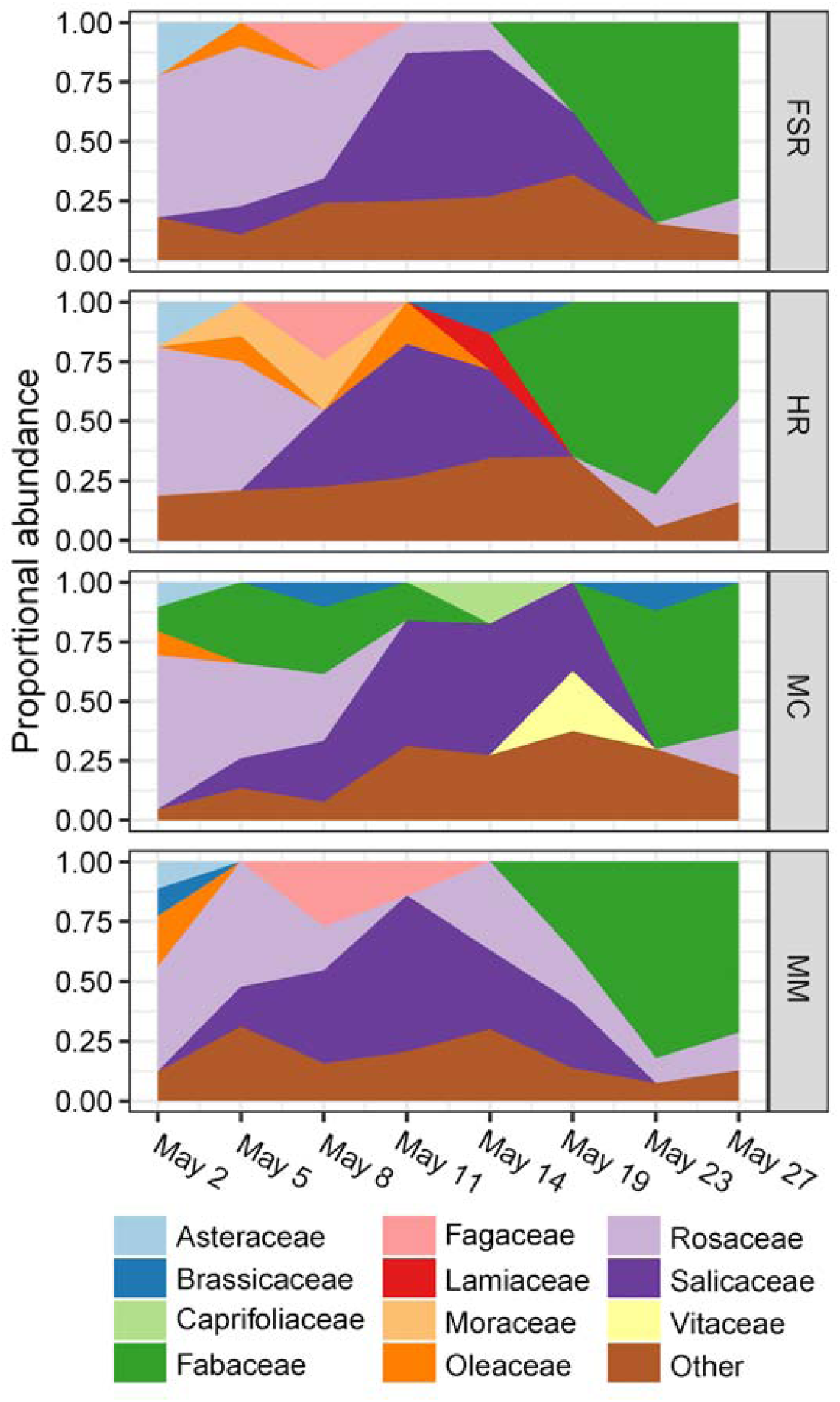
Time series plot of the metabarcoding median estimate of the proportional abundance of each plant family across the four sampling sites. Families occurring at lower than 10 percent abundance are not differentiated.

**Figure 4:**
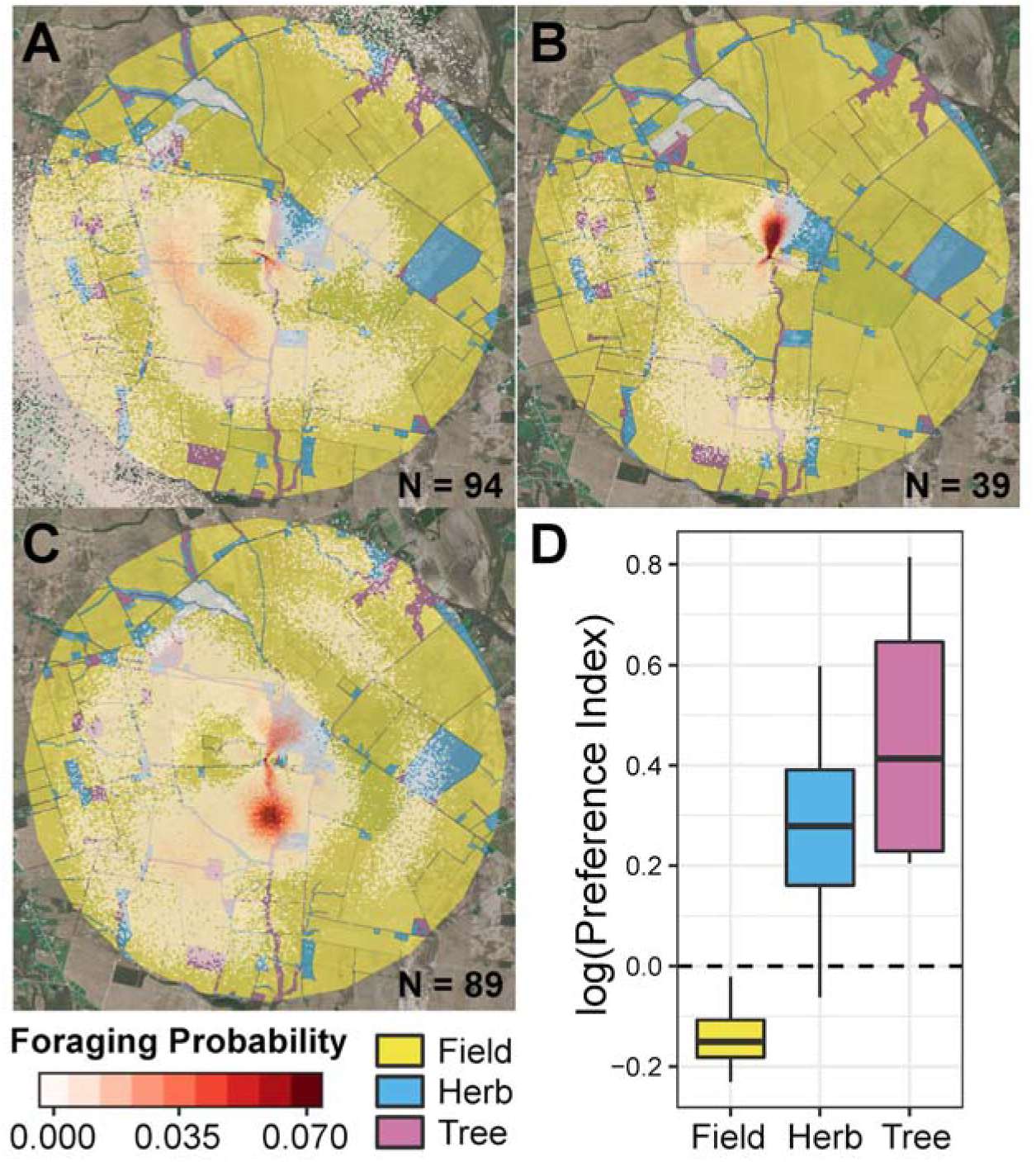
Honey bee spatial foraging patterns from May 2^nd^ to May 8^th^ (A), May 11^th^ to May 19^th^ (B) and May 23^rd^ to May 27^th^ (C) at one of the four sites. These sampling partitions represent the three major foraging periods observed in our data, dominated by Rosaceae, Salicaceae and Fabaceae, respectively. Boxplots of the log-transformed preference index across each of the three landcover types for all sites and all sampling dates (D). In total, 640 dances were analyzed for this work, with the sample size across sites ranging from 124 to 222 dances.

### Pollen foraging patterns

Our data indicate that three plant families, Rosaceae, Salicaceae and Fabaceae, comprised the majority of any given sample, accounting for a mean of 68.1 ± 3.0 (SE) percent of pollen abundance across all 32 samples. Results from the ITS2 locus, which displays greater resolution at lower taxonomic levels relative to other plant barcoding loci, led us to conclude that these family-level inferences likely represented *Prunus, Malus, Rubus, Salix, Trifolium* and *Cercis*.

### Waggle dance inferences

In analyzing the log-transformed preference index for each landcover type across all four sites, two-tailed t-tests suggested an overall preference for forested areas (mean log-transformed preference index: 0.4614; *P* = 0.0512) and an aversion to crop fields (mean log-transformed preference index: −0.1384; *P* = 0.0505). Though the mean preference index for non-crop herbaceous lands was positive, we did not observe a significant preference for this landcover class (mean log-transformed preference index: 0.2732; *P* = 0.1377). With respect to relative preferences, we found a significant difference in preference across landcover classes (one-way ANOVA: *P* = 0.01613). Specifically, forest areas were significantly preferred over crop fields (Tukey’s HSD test: *P* = 0.01444). Further, this effect did not appear to be driven by variation in the average distance of each landcover type from the hive, as we found no significant differences in this measurement (one-way ANOVA: *P* = 0.8943).

## Discussion

With our modified library preparation methods, we had three major goals: 1) obtain enough sequences per locus to accurately document the diversity of each sample, 2) obtain an even distribution of sequences per library, and 3) infer the taxonomic composition of our samples in a quantitatively representative manner. Considering past evaluations of the minimum number of analyzed pollen grains (Lau et al. 2017) and the sequencing depth needed to characterize the diversity of a typical sample of honey bee-collected pollen (Keller et al. 2015; Cornman et al. 2016), we are confident that our methods provided sufficient sequencing depth. Across all four loci, the minimum number of high quality Viridiplantae sequences generated for a sample was 38,486 and only 2 of 128 libraries contained fewer than 2,000 Viridiplantae sequences.

Despite adequate sequencing coverage, it is clear that our methods can be further optimized to yield a more even distribution of sequences per locus for each sample. This was an interesting outcome considering that we mixed our marker libraries on an equimolar basis before sequencing and may be explained by variation in amplicon clustering efficiency across loci on the Illumina MiSeq flow cell. Such sequence clustering variation is known to occur on the basis of template length (Illumina Inc. 2016). Given significant differences in the number of sequences per locus obtained, future studies implementing these methods would benefit from the addition of fewer ITS2, *trnL* and *trnH* products and more *rbcL* products during the final pooling of libraries. Additionally, investing in longer sequencing length would likely improve the taxonomic resolution achieved with the *rbcL* and *trnH* markers.

With respect to the quantitative capacities of pollen metabarcoding, numerous conflicting conclusions exist within the literature. While some authors conclude that molecular pollen identification methods can be relatively quantitative if interpreted appropriately (Hawkins et al. 2015; Kraaijeveld et al. 2015; Richardson et al. 2015a), others maintain that pollen metabarcoding data yield poor quality quantitative results (Bell et al. 2017; Bell et al. 2018). Our data indicate that, while all metabarcoding loci have some degree of bias, plastid loci produce data that is more quantitative, at least at the family rank and for the taxonomic groups assessed here. Further, the use of four metabarcoding loci and a median-based approach enables the estimation of pollen type abundance with reasonable quantitative accuracy when compared to microscopic analysis. As discussed in Richardson et al. (2015a), the use of multiple loci along with a consensus-filtered, median or average-based approach exhibits promise in terms of limiting false discoveries while increasing the scope of detectable taxa and increasing the quantitative utility of the resulting data. We contend that estimating the median, as opposed the to the mean, is ideal for this approach in order to reduce the influence of statistical outliers.

Alternatively, when considering the results from each locus individually, studies relying on a single marker and primer set to characterize diverse pollen samples almost certainly exhibit deficiencies with respect to taxonomic scope of detection and relative quantification, especially when a ribosomal locus is targeted. Poor results are expected for such loci considering that angiosperms are known to exhibit variations as large as 19-fold and 173-fold in ploidy and ribosomal copy number, respectively (Prokopowich et al. 2003; Murray et al. 2005). While several research groups contend that individual ribosomal loci alone are sufficient for quantitative inference (Keller et al. 2015; Pornon et al. 2016; Smart et al. 2017), a clear consensus of evidence challenges this assumption.

While the *trnL* locus used here appeared most quantitatively useful, the severe limitations of this fragment and primer set make it impractical for a single locus approach. Even though this short locus, approximately 160 bp in length, was the only locus to be sequenced in its entirety and efficiently mate-paired in this study, it exhibited poor resolution and could only be used for identification beyond the family level extremely rarely (23 percent of sequences identified to genus with an estimated false discovery rate of 17.8 percent). While additional sequencing length may result in fewer false discoveries and greater resolution for longer markers like *rbcL* and the *trnL* (UAA) fragment (Taberlet et al. 2007), it would not improve the results obtained with this section of *trnL*. It is also important to note that *trnL* libraries were more deeply sequenced by a large margin relative to *rbcL*, which may partially account for the increased quantitative performance of *trnL* (Smith and Peay 2014). Further, the relatively low proportion of sequences assigned to family for *rbcL* and ITS2, 78 and 70 percent, may have negatively affected the regression statistics of these markers relative to *trnL* and *trnH*, for which 97 and 99 percent of sequences were assigned to family. With respect to *trnL* and all loci analyzed, it should be considered that the quantitative evaluations presented here, as well as those of Richardson et al. (2015a), are limited to the taxa of early spring honey bee foraging in central Ohio, USA. Considering that different loci and primer sets exhibit variable, taxon-specific biases, *trnL* may perform differently in terms of quantitative reliability on alternate groups of plant taxa.

Dance analysis indicated a foraging preference for forested areas, particularly relative to crop fields. This is consistent with many studies that have observed a major role of forest and forest edge in provisioning honey bees, particularly in agricultural landscapes (Sande et al. 2009; Odoux et al. 2012; Donkersley et al. 2014; Richardson et al. 2015a; Requier et al. 2015). While previous work has found a negative correlation between forest land cover and honey bee productivity in Ohio (Sponsler and Johnson 2015), this apparent inconsistency could be explained by a positive effect of forest edge within an agricultural matrix and a negative effect of unbroken canopy in a forested matrix. This interpretation is supported by the predominance in our samples of *Salix* and rosaceous trees, which are characteristically forest edge, forest understory, and forested waterway flora. Importantly, though, considering that honey bees forage for nectar in addition to pollen, we are unable to precisely infer the degree to which observed spatial foraging patterns reflect pollen foraging. Thus, future studies of this nature would benefit from simultaneously conducting honey pollen analysis in addition to corbicular pollen analysis and waggle dance inference. Lastly, the present study was carried out in spring, when the majority of trees flower; if the dance analysis were repeated later in the year, outside the flowering period of major tree species, we would predict an aversion to forested areas and a preference for weedy herbaceous plants (Sponsler et al. 2017).

## Acknowledgements

The authors thank J. Bengtsson-Palme, D. Denlinger and K. Goodell for laboratory space and advice; I. Barnes and P. Young for access to bees and apiary sites; N. Douridas for assistance at the Molly Caren Agricultural Center; and field assistance from A. Sankey, N. Riusech, and M. Blackson. This work was supported by a Project Apis m. - Costco Honey Bee Biology Fellowship to RTR, a Pollinator Partnership Corn Dust Research Consortium grant to RMJ and support provided by state and federal funds appropriated to The Ohio State University, Ohio Agricultural Research and Development Center (OHO01277).

## Author Contributions

RTR, RMJ, CHL and DBS conceived and designed the study. HRC, CHL, SS, RTR, and DBS conducted the research. RTR, EGM, SS, RMJ and LEH analyzed the data. RTR wrote the manuscript and all authors helped edit the manuscript.

## Data Accessibility

Sequencing results produced in this work have been deposited to the NCBI Sequence Read Archive under BioProject PRJNA489437. The command line arguments, Python code and Metaxa2 trained databases used during data processing and analysis can be found at https://github.com/RTRichar/QuantitativePollenMetabarcoding.

## Supplemental Files

Supplemental File S1: Spreadsheet containing primer sequences, PCR conditions and the sample dual-indexing design used in this study.

